# Parametric cognitive load reveals hidden costs in the neural processing of perfectly intelligible degraded speech

**DOI:** 10.1101/2020.10.02.324509

**Authors:** Harrison Ritz, Conor Wild, Ingrid Johnsrude

## Abstract

Speech is often degraded by environmental noise or hearing impairment. People can compensate for degradation, but this requires cognitive effort. Previous research has identified frontotemporal networks involved in effortful perception, but materials in these works were also less intelligible, and so it is not clear whether activity reflected effort or intelligibility differences. We used functional magnetic resonance imaging to assess the degree to which spoken sentences were processed under distraction, and whether this depended on speech quality even when intelligibility of degraded speech was matched to that of clear speech (i.e., 100%). On each trial, participants either attended to a sentence, or to a concurrent multiple object tracking (MOT) task that imposed parametric cognitive load. Activity in bilateral anterior insula reflected task demands: during the MOT task, activity increased as cognitive load increased, and during speech listening, activity increased as speech became more degraded. In marked contrast, activity in bilateral anterior temporal cortex was speech-selective, and gated by attention when speech was degraded. In this region, performance of the MOT task with a trivial load blocked processing of degraded speech whereas processing of clear speech was unaffected. As load increased, responses to clear speech in these areas declined, consistent with reduced capacity to process it. This result dissociates cognitive control from speech processing: substantially less cognitive control is required to process clear speech than is required to understand even very mildly degraded, 100% intelligible, speech. Perceptual and control systems clearly interact dynamically during real-world speech comprehension.

**Significance Statement:** Speech is often perfectly intelligible even when degraded, e.g., by background sound, phone transmission, or hearing loss. How does degradation alter cognitive demands? Here, we use fMRI to demonstrate a novel and critical role for cognitive control in the processing of mildly degraded but perfectly intelligible speech. We compare speech that is matched for intelligibility but differs in putative control demands, dissociating cognitive control from speech processing. We also impose a parametric cognitive load during perception, dissociating processes that depend on tasks from those that depend on available capacity. Our findings distinguish between frontal and temporal contributions to speech perception and reveal a hidden cost to processing mildly degraded speech, underscoring the importance of cognitive control for everyday speech comprehension.

## Introduction

In perfect listening conditions, the comprehension of speech is seemingly effortless for healthy young people. However, everyday listening conditions are rarely as good as in the laboratory, and speech understanding is often compromised by noisy environments, low-fidelity digital communication, and hearing impairment. Listeners must exert cognitive control to understand markedly degraded speech (Broadbent, 1958; Eckert et al., 2016; Fedorenko, 2014; Heald & Nusbaum, 2014; Johnsrude & Rodd, 2016; Pichora-Fuller et al., 2016; Rouault & Koechlin, 2018; Vaden et al., 2013). However, what about very mildly degraded, perfectly intelligible speech? Does this also require attention and cognitive control, and if so, how much? A powerful method for quantifying control demands is to measure how processing of speech changes with declining speech quality, and under distraction. Neuroimaging experiments have reveal that cingulo-opercular regions associated with cognitive control (Shenhav et al., 2013) and temporal regions associated with high-level speech perception (Hickok & Poeppel, 2007) are sensitive to speech intelligibility (Davis & Johnsrude, 2003; Eckert et al., 2016), lose speech sensitivity during distracting tasks (Sabri et al., 2008; Wild et al., 2012), and that activity in these regions reflects perceptual accuracy (Wild et al., 2012; Vaden et al., 2013, 2015, 2016).

The existing body of research generally supports a role for domain-general control networks in degraded speech perception, however this work has been limited in its ability to parcellate regions into those that are speech selective, and those that respond in a domain-general fashion to all task demands. In a previous neuroimaging experiment, we found a set of frontal and temporal regions in which activity correlated with intelligibility when participants attended to speech, but not when they attended to either visual or auditory distractor tasks (Wild et al., 2012). In this study, the clear and degraded speech were not matched on intelligibility, limiting our ability to dissociate general and specific contributions to speech perception. For example, a *domain*-*general* region that monitors or controls task performance would appear sensitive to speech intelligibility during comprehension tasks, but only because intelligibility is strongly correlated with accuracy. In contrast, responses in a *domain*-*specific* region involved in effortful speech processing would reflect speech, regardless of task relevance, as long as cognitive resources are available. These two functions are likely to be organized hierarchically, with domain-general control processes in inferior frontal regions, and speech-selective processing in temporal regions of the frontotemporal language processing system (Davis & Johnsrude, 2003; Evans & Davis, 2015; Hickok & Poeppel, 2007). In the current study, we compare perception of clearly spoken sentences with perception of sentences matched for intelligibility (near-perfect word report accuracy), and sentences with only slightly lower intelligibility (>90% word report accuracy), allowing us to dissociate intelligibility from putative control demands.

As in our previous experiment (Wild et al., 2012), we measured speech processing when listeners are either attending to speech or when they are performing a distracting task. In order to better understand the tradeoffs in resource allocation between these two concurrent tasks, we parametrically varied cognitive load, and compared BOLD responses to intelligibility matched clear and degraded speech under these different levels. This novel parametric manipulation distinguishes processes that depend on the relevance of speech for the current task (*task*-*dependent* control) from the amount of control that is available to aid perception (*load*-*dependent* control). This parametric approach can help identify domain-general processes (e.g., monitoring of task-relevant accuracy), and can clarify the role of control in domain-specific processes (e.g., identifying when speech processing has a graded vs all-or-none dependence on cognitive load).

We demonstrate that the focus of attention, whether individuals were listening to speech or doing multiple object tracking, had a strikingly different effects on neural response to clear and degraded speech in high-level speech regions. Whereas the responses in anterior insulae were consistent with domain-general performance monitoring, anterior temporal cortex was selectively recruited for speech perception, with a strikingly different response profile for clear and intelligibility-matched degraded speech under parametric cognitive load. These results reveal the division of labor within a classical fronto-temporal speech network, where cognitive control is required, and enhances speech perception, in challenging listening conditions.

## Methods

### Participants

Twenty-six individuals (15 females; M_age_ = 21.5, SD_age_ = 3.86) participated in this experiment after providing informed consent in accordance with the research ethics board at the University of Western Ontario. Participants were right-handed, native English speakers, with self-reported normal (or corrected-to-normal) vision and self-reported normal hearing. Two participants were removed before analysis, due to dislodged earbuds or excessive movement during scanning, leaving 24 participants for the subsequent analyses.

### Experimental Design

On every trial, participants both heard a sentence and saw moving dots (See Figure 1). At the beginning of each trial, we instructed participants to either attend to the speech (‘LISTEN’), or to perform a visual tracking task (‘TRACK’). Across trials, we manipulated which task participants performed (2 levels), the clarity of speech that participants heard (3 levels), and the number of dots that participants saw on their screen (4 levels), generating 24 factorial conditions. Participants experienced 3 trials from each condition in each of the 3 scanning runs, for a total of 216 experimental trials. Participants also experienced two types of control trial: 24 silent, fixation-only trials and 24 LISTEN trials with rotated NV speech (see below), distributed equally across the three runs. We block-randomized conditions within each scanner run to minimize the effect of low-frequency drift.

**Figure 1.**
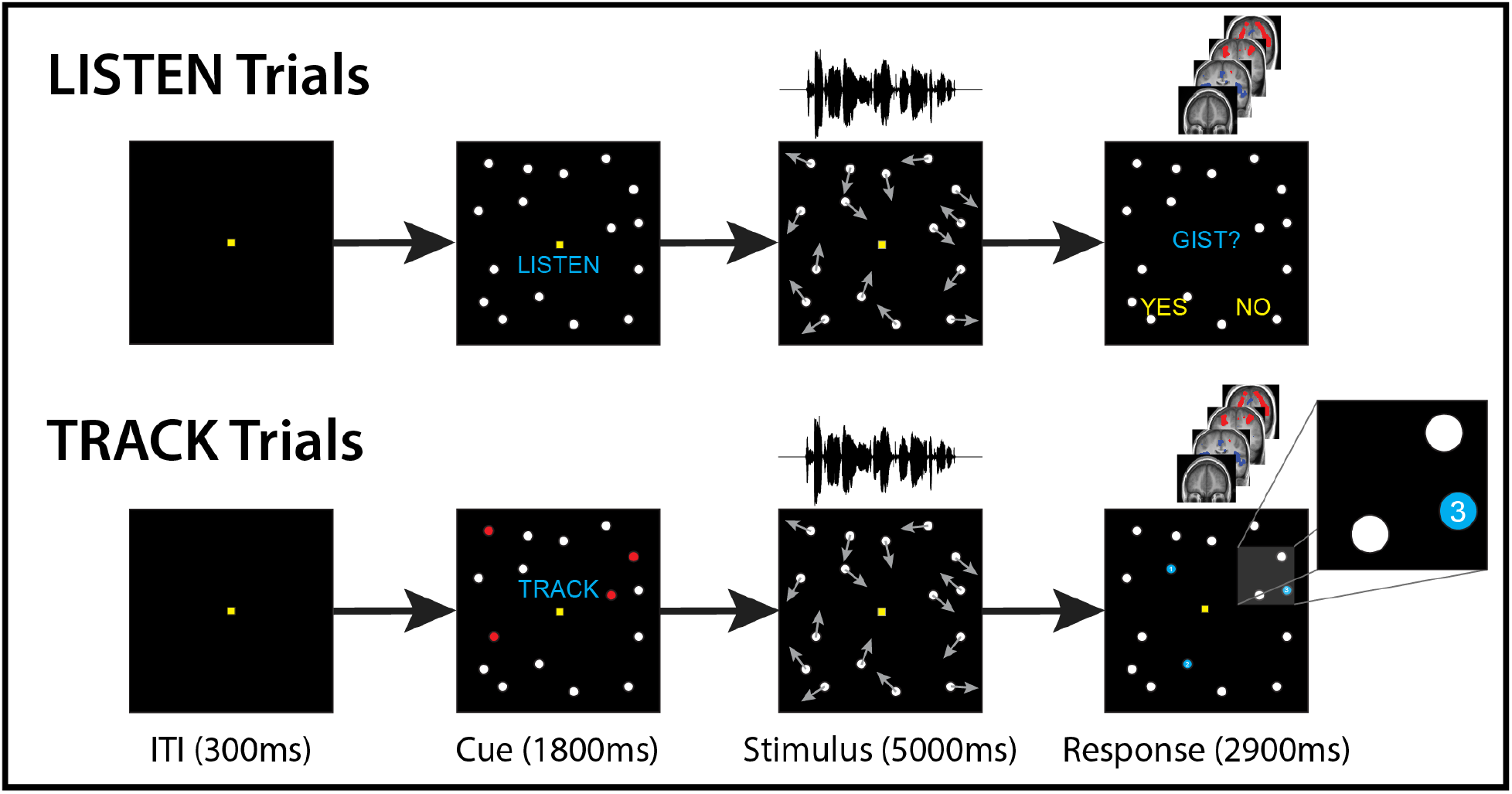
Trial Timecourse. At the beginning of each trial, participants were first cued to focus on speech (LISTEN) or focus on tracking (TRACK). They then both heard speech and saw moving dots, making a response during the wholebrain fMRI acquisition (occurring 4 seconds after stimulus midpoint). Speech stimuli were ordinary sentences (e.g., ‘Her handwriting was very difficult to read’) that were either clear (undistorted), 12-band noise-vocoded, or 6-band noise-vocoded, and during LISTEN trials participants reported whether they understood the ‘gist’ of each sentence. During tracking, participants tracked 1, 3, 4, or 6 moving dots among 12 distractors, and then reported which queried dot had been a member of the tracked set.

### Speech Task (LISTEN)

Due to a technical error, the comprehension and tracking data during scanning were lost for 2 participants, leaving 22 participants for behavioral analyses.

We used the same materials used in (Wild et al., 2012): 216 everyday sentences, all recorded by the same female speaker of Canadian English (e.g., ‘His handwriting was very difficult to read’). Stimuli were presented diotically via foam-tipped insert earphones (Sensimetrics, Belmont, USA) at a comfortable listening level. The sentences were 6-13 words long; 1.2-4.7 seconds in duration; and were split into six lists that were closely matched on the number of words, the sentence duration, and the summed word frequency (Thorndike and Lorge written frequency). These lists were assigned to the six Speech × Task conditions, counterbalanced across participants.

The clarity of the speech stimuli was manipulated using noise vocoding (Shannon et al., 1995). Each speech signal was filtered into logarithmically spaced frequency bands, with boundaries chosen to be equally spaced along the basilar membrane (Greenwood, 1990). The amplitude envelope within each frequency band was extracted and convolved with white noise that was band-limited to the same frequency range. Previous work has found that intelligibility depends on the number of bands (Davis & Johnsrude, 2003; Shannon et al., 1995). In this experiment, we used highly intelligible noise-vocoded stimuli, filtered with 12 (NV12) and 6 (NV6) bands, as well as Clear (un-manipulated) speech. Piloting and previous experiments have determined that people can accurately report nearly 100% of the words from both Clear and NV12 sentences, whereas word-report of NV6 speech is poorer, but still greater than 90% (see Figure 2A). Unintelligible, spectro-temporally matched, control stimuli were generated by “spectral rotation”: during the vocoding process, we permuted the assignment of speech envelopes to their noise envelopes (i.e., randomized over frequency bands; (Blesser, 1972)).

**Figure 2.**
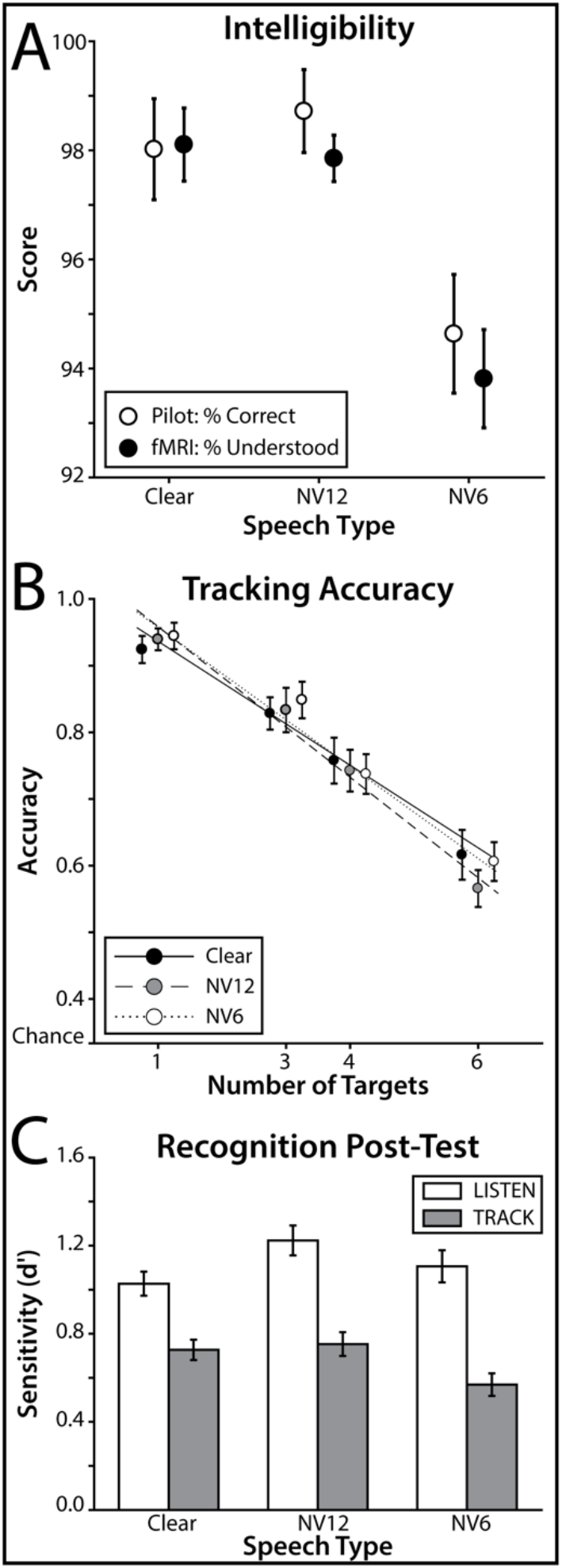
Behavioral Results. **A**: Intelligibility. Intelligibility scores across speech types were similar whether measured as objective word report accuracy (behavioral pilot; *n* = 12) or as subjective gist report (scanner experiment; *n* = 22). **B**: Tracking Accuracy. When participants tracked more targets, their tracking accuracy declined. Participants’ accuracy remained above chance (33%) at all levels of tracking load. **C**: Recognition Post-Test. After the main experiment, participants performed a surprise memory test for the speech stimuli, deciding whether written probes had been heard previously or were novel. Memory sensitivity was quantified with d’, comparing hit and false alarm rates. All error bars indicate within-participant SEM (Morey, 2008).

Volumes were collected using a sparse acquisition protocol (Hall et al., 1999), in which our speech stimuli were presented during the silent period (9 seconds) between scans. The onset of each scan began 4 seconds after the midpoint of each sentence and tracking task, sampling the hemodynamic response near its peak amplitude. On LISTEN trials, participants had 2.8 seconds near the end of the 9-sec silent period to indicate with a yes/no keypress (dominant hand) whether they had understood the gist of the sentence (see Figure 1).

### Multiple Object Tracking Task (TRACK)

Between 13 and 18 dots were on the screen throughout every trial, regardless of the task. All dots had a diameter of ~1 degree of visual angle and were shown against a black screen spanning ~ 20 × 20 degrees. Dots were stationary for 1.8 seconds, and then moved pseudorandomly around the screen at an approximate speed of 1.8 degrees per second, with dots repelling 180 degrees away from other dots or the edge of the screen at a 0.5-degree proximity.

On TRACK trials, participants tracked a subset of the moving dots (multiple object tracking, MOT; Pylyshyn & Storm, 1988). On these trials, 1, 3, 4, or 6 target dots were highlighted in red for 1.8 seconds before movement. Participants were instructed to keep their gaze on a fixation cue in the center of the screen and track these dots covertly. After 5 seconds of tracking, the dots froze in place, and three dots (one randomly selected target and two foils) were highlighted in blue and labelled ‘1’, ‘2’, and ‘3’. Participants had 2.8 seconds to indicate with a 3-alternative keypress which of the numbered dots was a target, without feedback (see Figure 1).

### Pre-Training and Memory Post-Test

Prior to the scanning session, participants practiced both the speech and tracking tasks. First, participants were familiarized with NV speech, in order to bring their comprehension performance to asymptote (Davis et al., 2005). Over 24 trials, participants heard a noise-vocoded sentence, indicated whether they had understood the gist of the sentence, and then received feedback by hearing the vocoded sentence again while also reading it on the screen (following the recommendations from Davis et al., 2005, Experiment 3). MOT training proceeded over 24 trials. On the first 12, the number of targets began at 1 and increased (to 3, 4 and 6) after each correct tracking response, and decreased after each incorrect response. On the latter 12 trials, the number of targets on each trial was randomly selected (from 1, 3, 4 or 6).

After the scanning session, we tested participants on their recognition memory for the sentences they had heard. On each trial, participants saw a written sentence on a computer screen, and indicated with a keypress whether they remembered this sentence from the experiment (‘OLD’), or whether it was new (‘NEW’). Participants were tested on all 216 sentences from the experiment, along with 108 foil sentences. Foil sentences differed from target sentences in both their topic and their content words. During the scanning session, participants were unaware that memory would be tested, ensuring incidental memory encoding.

### fMRI acquisition

Images were acquired on the 3.0T Siemens Prisma MRI system at the University of Western Ontario. T1-weighted structural images were collected at the beginning of each session using a single-shot EPI (FoV: 256mm^2^; resolution: 1mm isotropic; slice thickness: 1mm with 50% gap; TE: 2.98ms; TR: 2300ms; flip angle: 9°). T2*-weighted functional volumes were acquired across the whole brain using a 4-factor interleaved multi-band gradient EPI (FoV: 192mm^2^; resolution: 2.5mm isotropic; slice thickness: 2.5mm with 10% gap; 52 slices; TE: 30ms; TA: 1000ms; TR: 10sec; flip angle: 70°). Acquisition was transverse-oblique, angled away from the eyes.

### fMRI preprocessing and analysis

fMRI data were preprocessed and analyzed using SPM12 (Wellcome Centre for Neuroimaging, London, UK), following standard preprocessing steps including realignment, coregistration, and simultaneous segmentation and normalization to MNI (ICBM452) space. Normalization parameters were calculated from the structural image and applied to functional images coregistered to the mean of each run, resampling the images at 2mm^3^. The normalized images were spatially smoothed using a 3D Gaussian kernel with an 8mm FWHM.

Statistical parametric maps for each subject were estimated using a general linear model containing onset indicators for rotated speech and the six combinations of Speech (Clear, NV6, and NV12) by Task (LISTEN and TRACK) conditions. The model also included Load parametric modulators for the six speech × task conditions, based on the dots on the screen. For LISTEN trials, the parametric modulators only captured the number of dots on the screen (c.f., visual load), whereas for TRACK trials, these modulators also captured the effect of tracking load. These models also included run-specific modulators including the six spatial realignment parameters, as well as a run intercept and linear trend. Modulators were mean centered and not orthogonalized (allowing control modulators to compete for variance with task modulators). Due to the long TR (10 seconds; 9 second silent gap between successive scans) in our sparse acquisition design, we modelled trial activation using a finite-impulse response model without serial autocorrelations. Contrast maps for main effects and interactions were calculated at the subject level and tested against zero at the group level using a factorial partitioned-error repeated-measures ANOVA (Henson & Penny, 2003).

We analyzed participants’ behavior using custom MATLAB (R2018a) scripts and JASP (0.8.3) for ANOVA and Bayesian analyses (using the default Cauchy prior). Note that Bayes factors (BF_10_) less than 1/3 provide moderate evidence supporting the null hypothesis (e.g., that two groups are the same; see Jarosz & Wiley, 2014). Follow-up fMRI analyses were performed using MATLAB and JASP. For our follow-up interaction analyses, we utilized a second general linear model that included all twenty-four Speech × Task × Load conditions, along with our run-specific nuisance terms (see above). We followed-up omnibus ANOVAs with post-hoc t-tests, correcting for multiple comparisons with the Holm procedure for sequential tests. Brain-behavior relationships were crossvalidated by fitting a linear regression model to predict BOLD contrasts from behavior while holding-out one participant at a time, using this model to predict each held-out participant’s BOLD contrast from their behavior, and then correlating the predicted and observed BOLD contrasts.

## Results

### Task overview

At the beginning of each trial of the fMRI session, participants were instructed to perform one of two tasks (see Figure 1). During LISTEN trials, they reported whether they understood the ‘gist’ of a sentence that was either not degraded (Clear), degraded but as intelligible as clear speech (NV12), or degraded below the intelligibility of clear speech (NV6; still over 90% intelligible). On a subset of LISTEN trials participants instead heard unintelligible speech (Rotated), a spectrotemporal-matched acoustic baseline for speech.

During TRACK trials, participants performed MOT, tracking either 1, 3, 4, or 6 pseudorandomly moving dots among 12 distractors, and then reporting which out of three highlighted dots had been a member of the tracked set (33% chance rate). Participants saw different numbers of moving dots during LISTEN trials and heard different kinds of speech during TRACK trials, in a fully crossed factorial design consisting of Task (2 levels), Speech Type (3 levels), and number of dots (4 levels). After the scanning session, participants performed a recognition memory test for sentences presented during the scan session (visually presented one at a time), as a secondary measure of their speech comprehension.

### Task Performance

During LISTEN trials, participants reported whether they understood the gist of each sentence. Participants reported understanding almost all of the intelligible speech trials (Clear: 98.1%; NV12: 97.9%; NV6: 93.8%), and almost none of the Rotated trials (5.3%). These scores were similar to the word-report accuracy collected from a separate group of pilot participants (all BF_10_ ≤ 0.5; see Figure 2A). Gist scores differed among intelligible speech types (*F*_(1.26, 26.6)_ = 12.1, *p* < .001, η^2^= .365). Whereas Clear and NV12 did not differ (*p*_Holm_ = .648, BF_10_ = 0.246), gist scores were higher for both Clear and NV12 compared to NV6 (Clear: *t*_(21)_ = 3.44, p_Holm_ = .005; NV12: *t*_(21)_ = 3.98, *p*_Holm_ = .002).

During TRACK trials, participants tracked 1, 3, 4, or 6 moving dots and then selected the member of the tracked set with a three-alternative forced choice. Participants’ tracking accuracy linearly decreased as load increased (logistic mixed-effects regression: β = −0.44, *t*_(21)_ = −11.0, *p* < .001), from 94% accuracy for 1 dot to 60% accuracy for 6 dots. Participants consistently performed above chance (33%), even at the highest level of Load (6 dots: *t*_(21)_ = 11.1, *p* < .001; see Figure 2B).

Sensitivity scores (d’) indexing how well participants could distinguish sentences heard during the experiment from foils during the post-scan recognition memory test are depicted in Figure 2C. Sentences from all conditions were remembered better than chance (one-sample t-tests against 0: all *p*_Holm_ < .001). We ran a 3 (Speech) × 2 (Task) repeated-measures ANOVA on d’ scores, finding that participants remembered sentences better during LISTEN than TRACK trials (*F*_(1, 23)_ = 81.0, *p* < .001, η^2^ = .779).

Recognition memory also depended on Speech type (*F*_(1.94, 44.7)_ = 5.64, p = .007, η^2^ = .197): memory for NV12 speech was significantly better than for NV6 (*t*_(23)_ = 3.17, *p*_Holm_ = .008) and marginally better than for Clear speech (*t*_(23)_ = 2.26, *p* = .057). The interaction between Task and Speech Type was only marginally significant (*F*_(1.99, 46.0)_ = 2.84, *p* = .069, η2 = .110), suggesting that memory for NV12 speech may have benefited from the LISTEN task more than Clear speech (*t*_(23)_ = 2.31, *p*_Holm_ = .076). This interaction matches our previous observations of stronger memory performance for degraded than clear speech when these are both attended (Wild et al., 2012). These memory results suggest that performance of the MOT task disrupted speech processing, and that attention to mildly degraded sentences enhances processing.

### Task-specific neural responses

Participants appeared to orient their attention depending on the task cue (Figure 3). Consistent with previous studies, LISTEN trials elicited greater activity across temporal and lateral prefrontal cortices (Davis & Johnsrude, 2003; Scott et al., 2000), whereas TRACK trials elicited greater activity in posterior parietal and superior frontal cortices (Culham et al., 2001; Howe et al., 2009).

**Figure 3.**
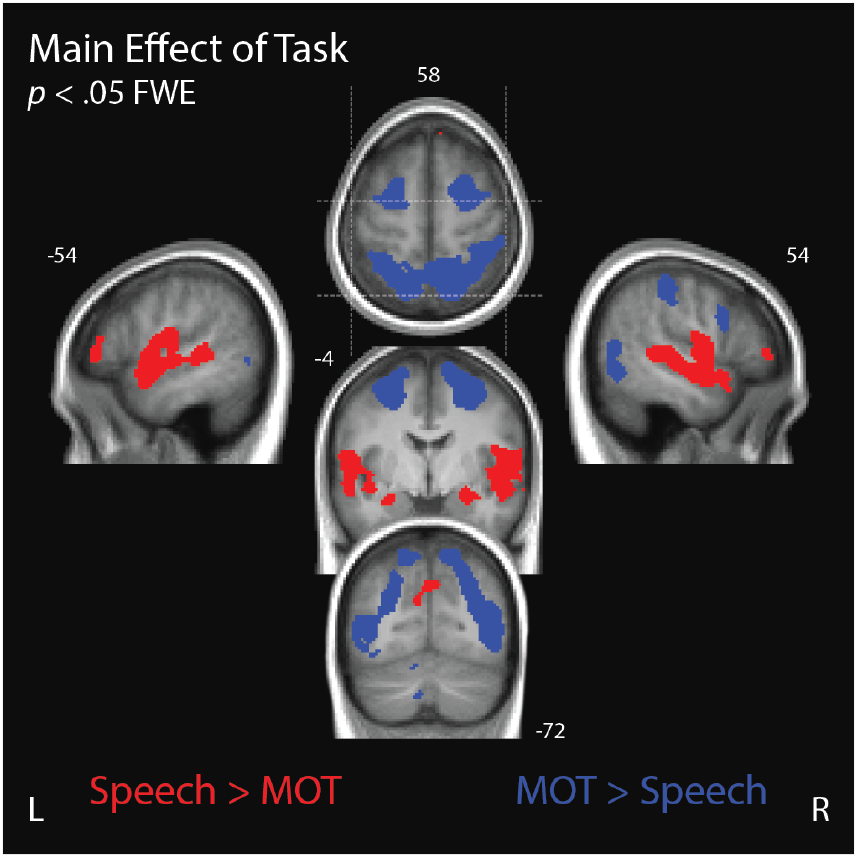
Main effect of Task. Voxels that exhibited a significant main effect of Task were colored according to whether they exhibited a greater response to LISTEN than TRACK, or vice versa (*p* < .05, whole-brain FWE). Activation is plotted on the mean participant T1-weighted structural MR image, with dashed lines on the axial slice indicating the location of the sagittal and coronal slices. See supplementary materials for coordinate table.

We tested the simple main effect of Speech Type during LISTEN trials only, as we hypothesized that speech processing would depend on attention (see Figure 4A). Comparing the activity elicited by Clear, NV12, NV6, and Rotated speech during LISTEN trials, we observed a simple main effect of Speech Type across temporal and cingulo-opercular cortices. Temporal lobe voxels appeared to be sensitive to the intelligibility of speech, exhibiting progressively greater activity as gist report accuracy increased across the four speech types (green voxels; Davis & Johnsrude, 2003; Wild et al., 2012). In contrast, cingulo-opercular voxels exhibited greater activity for NV6 speech than for clear and NV12 speech (blue voxels), consistent with these regions responding more when stimuli are degraded (Eckert et al., 2016; Wild et al., 2012). These hypothesis-driven contrasts were not exhaustive, and some regions showed a main effect of speech with a different pattern of activation (white voxels).

**Figure 4.**
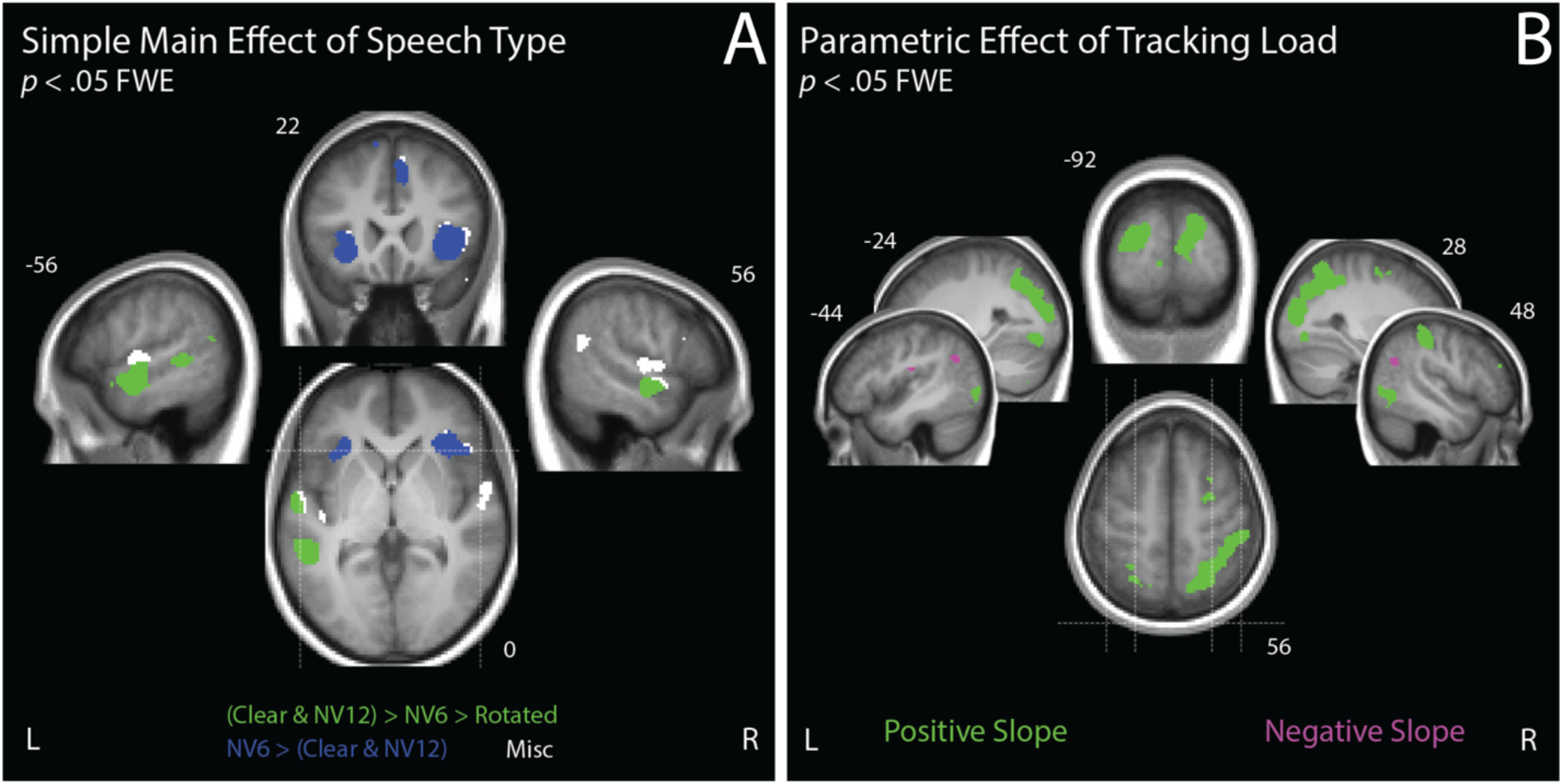
Task-Specific Simple Main Effects. **A:** Simple Main Effect of Speech Type. Voxels that exhibited a significant simple main effect of Speech Type (Clear, NV12, NV6, or Rotated) during LISTEN are colored according to hypothesized contrasts (Wild et al., 2012). Green voxels indicate a greater response for more intelligible speech and blue voxels indicate a greater response for NV6 compared to more intelligible Clear and NV12 speech. (White voxels exhibited any simple main effect pattern not captured by these contrasts.) **B:** Parametric Effect of Tracking Load. Voxels that exhibited a significant parametric effect of the number of dots tracked during TRACK are colored green if they show a positive relationship, and magenta if they show a negative relationship. In both images, activation is shown on the mean participant T1-weighted structural MR image, and dashed lines on the axial slice indicate the location of the sagittal and coronal slices. See supplementary materials for coordinate table.

Despite the highly similar intelligibility of Clear and NV12, our neural measures distinguished these speech types. Contrasting Clear vs NV12 during LISTEN revealed a significant peak in the left STG (*F*_(1, 23)_ = 80.46, *p* < .001, whole-brain FWE) and a marginally significant peak in the right STG (*F*_(1, 23)_ = 40.91, *p* = .069). These clusters partially overlapped with intelligibility-sensitive regions. Both STG regions were more sensitive to Clear than to NV12 speech. No voxels exhibited a significantly stronger response to NV12 than to Clear.

Finally, we tested for the simple parametric effect of tracking load during TRACK. In many of the regions that were more active for TRACK than LISTEN (main effect of task), BOLD activity was positively correlated with tracking load (see Figure 4B; green voxels), consistent with previous reports (Bettencourt, 2010; Culham et al., 1998, 2001; Howe et al., 2009; Jovicich et al., 2001; Tomasi et al., 2004). We also observed negative correlations with tracking load in the left supramarginal gyrus and angular gyri bilaterally (magenta voxels).

### Domain-general response in anterior insulae

Our primary hypotheses concern the degree to which speech processing requires attention under different levels of degradation. Accordingly, we tested our 2- and 3-way interactions within a large speech-sensitive mask based on our previous investigation (Wild et al., 2012), which was fully independent of the current experiment. We defined our mask as voxels exhibiting either a significant main effect of Speech Type or a Speech Type × Task interaction in this previous experiment (see Figures 4 and 5 from Wild et al., 2012).

**Figure 5.**
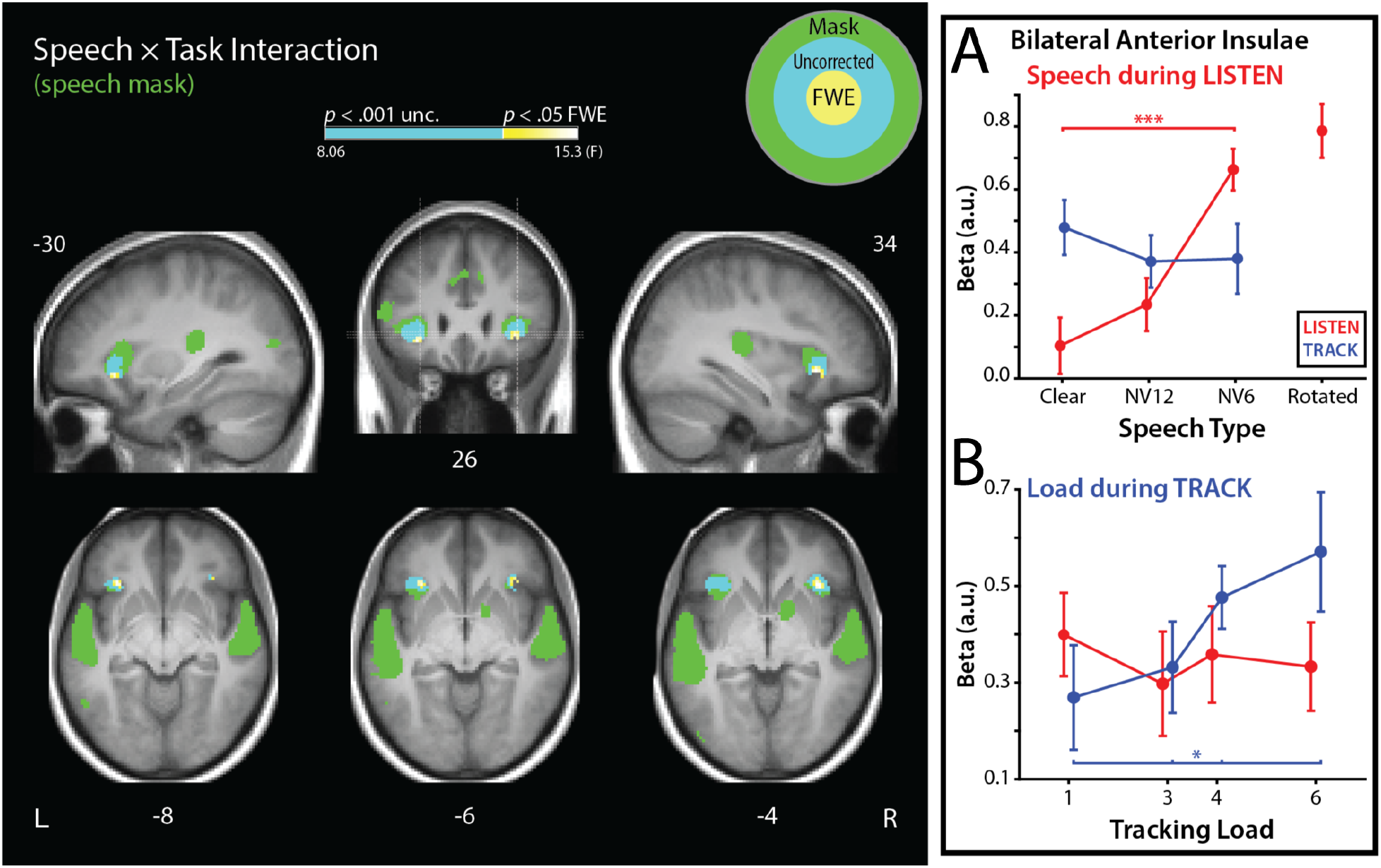
Speech × Task Interaction. Analyses were performed within an independent mask of speech-sensitive cortex (green; see text). Cyan voxels exhibited an interaction between Speech Type and Task at an uncorrected threshold (p < .001). Voxels that exhibited a significant interaction at a corrected threshold are indicated with a heat map corresponding to their F-statistic (*p* < .05, within-mask FWE). **A**: Parameter estimates extracted from above-threshold voxels show a significant simple main effect of Speech Type only during LISTEN (red). **B**: A post-hoc analysis found a significant positive parametric effect of Load only during TRACK (blue). Error bars indicate SEM adjusted for within-subject measurements (Morey, 2008). Activation is plotted on the mean participant T1-weighted structural MR image, and dashed lines on the coronal slice indicate the location of the sagittal and axial slices. See supplementary materials for coordinate table.

We observed a significant interaction between Task (LISTEN and TRACK) and Speech Type (Clear, NV12, and NV6) in the anterior insulae bilaterally, consistent with our previous experiment (Wild et al. 2012b; see Figure 5). To compare the response profiles across hemispheres, we ran a Hemisphere × Speech Type × Task mixed ANOVA on the parameter estimates from these regions. The hemisphere factor did not influence our interaction effect (BF_10_ = .201), so we averaged parameter estimates across above-threshold voxels in this region across hemispheres.

In this insular region was a simple main effect of Speech Type during LISTEN (*F*_(1.87, 43.1)_ = 18.65, *p* < .001) that was not significant during TRACK (*F*_(1.53, 35.2)_ = 0.458, *p* = .585; BF_10_ = .172; see Figure 5A). During LISTEN, the anterior insulae’s response was greater for NV6 than Clear speech (*t*_(23)_ = 5.81, *p*_Holm_ < .001), and NV12 speech (t_(23)_ = 5.10, *p*_Holm_ < .001). Activation during LISTEN for Clear and NV12 speech did not differ (*p*_Holm_ = .229; BF_10_ = .423). This pattern of elevated activity for difficult-to-understand degraded speech (NV6), only when this speech is task-relevant, is consistent with the response profile observed in (Wild et al., 2012).

To further characterize the task-dependent role of the anterior insulae, we also tested whether the effect of tracking load was evident in these insular voxels (see Figure 5B). We found that the insular response linearly increased with Load during TRACK (*t*_(23)_ = 2.22, *p* = .036), with a stronger Load effect during TRACK than LISTEN (*t*_(23)_ = 2.55, *p* = .018). Together, these signals suggest that the insulae’s response reflected the performance of the currently attended task.

### Domain-specific response in anterior temporal cortex

Our analysis of primary interest examined whether there are speech-sensitive regions in which the effect Speech Type depends on the load during TRACK trials, and in particular whether this cognitive load dissociates processing of Clear speech from intelligibility-matched degraded speech (NV12). Using the same speech-sensitive mask as our Speech Type × Task analysis, we examined the interaction of Speech Type × Task on the parametric Load modulators (effectively examining the Speech × Task × Load interaction). We found that this interaction was significant in anterior portions of the superior temporal gyri bilaterally (aSTG; see Figure 6). As with the insulae, we found that this interaction was similar across hemispheres (BF_10_ = .301), so we averaged the parameter estimates across above-threshold voxels in both hemispheres.

**Figure 6.**
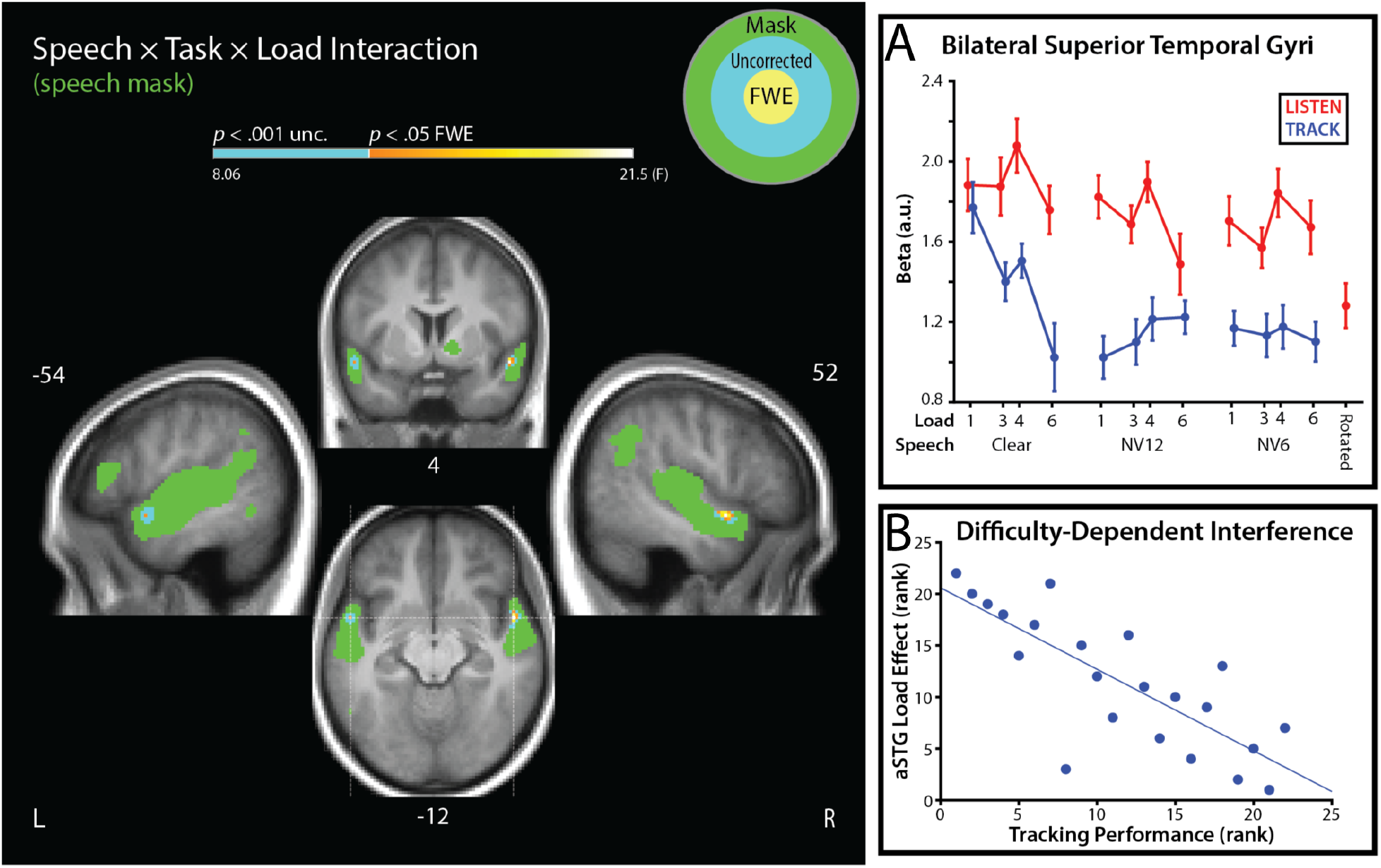
Speech × Task × Load Interaction. Analyses were performed within an independent mask of speech-sensitive cortex (green; see text). In cyan voxels, the slope relating BOLD activation to tracking load depended on both Task and Speech Type (*p* < .001, uncorrected). Voxels that exhibited a significant interaction at a corrected threshold are indicated with a heat map corresponding to their F-statistic (*p* < .05, within-mask FWE). **A**: Parameter estimates extracted from above-threshold voxels show a different load response for Clear and degraded speech during TRACK (blue), with degraded speech yielding activation in these regions at floor level (defined by the Rotated-speech point) at all tracking loads. In marked contrast, activity for Clear speech did not depend on task when Load was low (1-item MOT), but then linearly declined with increasing tracking load. **B**: Participants who had more difficulty with the tracking task (lower Accuracy / RT) had a stronger interaction between Speech Type and Load during TRACK. Error bars indicate within-subject SEM (Morey, 2008). Activation is plotted on the mean participant T1-weighted structural MR image, and dashed lines on the coronal slice indicate the location of the sagittal and axial slices. See supplementary materials for coordinate table.

During LISTEN, the effect of Load was not significant, nor was there a Load × Speech Type interaction (*F*_(2, 46)_ = 1.38, *p* = .267, BF_10_ = .278). This was expected, since Load predictors during LISTEN only indexed the number of (task-irrelevant) dots on the screen. In contrast, during TRACK, the parametric Load effect depended on Speech Type (*F*_(2, 46)_ = 12.13, *p* < .001; see Figure 6A). The Load effect was apparent for Clear speech, with activity decreasing as load increased beyond 1-item MOT. In contrast, for NV12 and NV6 speech, activity during TRACK was at floor even for 1-item MOT, eliciting a response no stronger than for unintelligible rotated speech (Load_Clear_ - Load_NV12_: *t*_(23)_ = −4.041, *p*_Holm_ < .001; Load_Clear_ - Load_NV6_: *t*_(23)_ = −2.92, *p*_Holm_ = .016). Across all of the Speech conditions in both tasks, only Clear speech during TRACK exhibited a significant effect of Load (Clear during TRACK: *t*_(23)_ = −3.20, *p*_bonferroni_ = .024; all other *p*_uncorrected_ ≥ .16 and BF_10_ ≤ .545).

Another way to compare our Speech conditions is to examine, within each Speech Type, the MOT load at which differences between tasks begin to arise. Within each Speech Type, therefore, we compared the response at each level of Load during TRACK to the overall response during LISTEN. When one target was being tracked (lowest load), the STG response for clear speech was similar between TRACK and LISTEN (*t*_(23)_ = −1.02, *p*_uncorrected_ = .32, BF_10_ = 0.344). In marked contrast, activity evoked by degraded speech depended strongly on Task: activity for both NV12 and NV6 was substantially lower during TRACK than LISTEN, even at the weakest level of Load (NV121-target: *t*_(23)_ = −6.07, *p*_Holm_ < .001; NV61-target: *t*_(23)_ = −5.76, *p*_Holm_ < .001). When tracking three or more objects, STG activity was always lower for TRACK than LISTEN, and did not differ among speech types (Effect of Speech Type when Load > 1: BF_10_ = 0.038).

Complementing our neural measures, we also examined whether individual differences in the strength of this Load by Speech Type interaction was correlated with participants’ task performance. We found that participants’ with a stronger aSTG Load effect during TRACK (Load_(NV12, NV6)_ - Load_Clear_) had worse average overall tracking accuracy (Spearman’s correlation: ρ_(20)_ = −.46, *p* = .032) and slower median reaction times (ρ_(20)_ = .52, *p* = .014). We validated the generalizability of these individual differences using a leave-one-out cross-validation procedure. A measure of processing efficiency (accuracy / RT) was strongly correlated with aSTG Load effects within-sample (ρ_(20)_ = −.79, *p* < .001; see Figure 6B), and regression predictions for held-out participants strongly correlated with their performance (ρ_(20)_ = .74, *p* < .001). Participants with stronger neural indicators of load-dependent interference on speech processing performed more poorly on the MOT task, suggesting that our aSTG neural measures reflect the subjective task demands.

In sum, the response to clear speech in anterior temporal cortex was similar regardless of the focus of attention when tracking was easy, but linearly declined to the same low level as for degraded speech with increasing tracking load. This neural index of interference was more severe for participants that were overall worse at the tracking task. The response profile for clear speech was fundamentally different from that for equally intelligible degraded speech, with activity for this degraded speech at the same level as unintelligible Rotated speech, even at weakest level of tracking load.

## Discussion

Intelligibility responses in the anterior portion of the ventral speech pathways depend on attention (Eckert et al., 2016; Sabri et al., 2008; Wild et al., 2012). In the current experiment, we found that these regions can be fractionated based on whether speech-sensitivity depends on the current task or the available processing capacity. Activity in the anterior insulae appeared to reflect the demands of the instructed task. This region responded more strongly to more degraded speech only when speech was task-relevant, and activity depended linearly on tracking load only during MOT (Figure 5). In contrast, sensitivity to speech in anterior temporal regions depended both on the type of speech and, for clear speech, on concurrent cognitive demands (Figure 6). This load-dependent response in bilateral temporal lobes strongly dissociated clear speech from intelligibility-matched degraded speech: clear speech was unaffected by the weakest level of distraction, at which the degraded speech response was already reduced to baseline. These observations functionally parcellate speech-sensitive cortex in the inferior frontal and superior temporal regions based on their relationship to cognitive control, demonstrating substantial costs of distraction under natural, perfectly intelligible, levels of speech degradation.

The anterior insulae play an important role in cognitive control (Bunge et al., 2002; Cieslik et al., 2015; Dosenbach et al., 2006; Duncan & Owen, 2000; Fedorenko et al., 2013; Shenhav et al., 2013), and may support performance monitoring (Lamichhane et al., 2016; Vaden et al., 2013; Wager et al., 2005) and/or orienting towards salient events (Craig & Craig, 2009; Klein et al., 2007; Seeley et al., 2007; Ullsperger et al., 2010). In this experiment, activity in the anterior insulae was sensitive only to the demands of the instructed task: stronger responses to degraded speech only during LISTEN (as in Wild et al., 2012b), and positive linear dependence on tracking load only during TRACK. During LISTEN, this region exhibited a similar response for clear and intelligibility-matched degraded speech, also consistent with a generic role for performance monitoring (Vaden et al., 2013, 2015, 2016).

In anterior temporal cortex, we found that speech sensitivity depends on the cognitive demands of a distracting task. When Clear speech was task-irrelevant, the aSTG response linearly declined as tracking load increased, with a stronger decline predicting poorer tracking performance. This decline may reflect a decreased availability of attention to enhance speech perception or active suppression of this region to reduce interference, with both accounts implying shared capacity for speech perception and MOT (Broadbent, 1958; Kahneman, 1973). MOT is a relatively simple task designed to isolate attentional processes that index object locations (Cavanagh & Alvarez, 2005; Pylyshyn & Storm, 1988; Scholl, 2009), with recent theoretical (Franconeri et al., 2010) and computational (Srivastava & Vul, 2016) models proposing that a critical function of MOT is protecting target indices from interference (i.e., from ‘swapping’ a target with a distractor; Pylyshyn, 2004). During speech perception, there may be analogous competition between phonological, lexical, and semantic candidates (e.g., multiple potential interpretations of a sound or word), which is exacerbated by degradation (Luce & Pisoni, 1998; Marslen-Wilson, 1987; Miller et al., 1951; Novick et al., 2005; Rodd et al., 2002; Spivey et al., 2005; Thompson-Schill et al., 1997; Zhuang et al., 2011). During both tasks, attention could plausibly be allocated in response to heightened uncertainty and competition (e.g., towards regions of target/distractor proximity in MOT, or proximal phonological candidates during speech), a core process in domain-general cognitive control (Berlyne, 1957; Miller & Cohen, 2001; Posner & Snyder, 1975).

When attention was on the MOT task the anterior temporal response to (task-irrelevant) intelligible degraded speech was eliminated, which contrasted markedly with the response during task-irrelevant clear speech. This profile may reflect ‘maxed-out’ processing capacity, or additional functions that are unavailable under distraction (e.g., functions that are goal-dependent). That processing capacity was entirely occupied by the MOT task is not likely, given that the response in anterior temporal regions to mildly degraded speech was at the baseline even when individuals were tracking a single object, which is a very modest level of load. Furthermore, the load effect was clearly evident for task-irrelevant clear speech, but not for degraded speech.

Instead, the processing of perfectly intelligible degraded speech in anterior temporal lobe regions appears to be gated by task goals. Consistent with this idea, activity in anterior insulae was determined by the demands of the attended task, plausibly in the service of top-down control over anterior temporal cortex (Novick et al., 2005; Wild et al., 2012; Eckert et al., 2016). The insulae and anterior temporal lobe share extensive anatomical connections via the uncinate fasciculus and extreme capsule (Kier et al., 2004; Petrides & Pandya, 1988, 2007; Romanski et al., 1999), which have long been thought to facilitate speech perception (Wernicke, 1908). Neuropsychological and neuroimaging evidence supports a role for this network in semantic processing (Dick & Tremblay, 2012; Hickok & Poeppel, 2007; Saur et al., 2008). For example, electrostimulation to extreme capsule fibers in the anterior insulae reliably induce ‘semantic paraphasias’, with patients replacing target words with semantically-related competitors (e.g., brush → comb; Duffau et al., 2005), a potential complement to the target-distractor swaps that characterize MOT performance (Pylyshyn, 2004; Franconeri et al., 2010; Srivastava & Vul, 2016). While these similarities are promising, further research is needed to fully characterize the neural interactions that support selective attention during speech perception.

Consistent with enhanced top-down control during degraded speech perception, recognition memory tended to be better for NV12 speech than Clear speech when it was the focus of attention (as in Wild et al., 2012; see also Hirshman & Mulligan, 1991; Nairne, 1988). However, these findings contrast with previous research that has documented poorer memory for degraded speech (Murphy et al., 2000; Pichora-Fuller et al., 1995; Rabbitt, 1966; Surprenant et al., 1999). In many of these previous experiments, stimuli lacked the contextual constraints of full sentences (Murphy et al., 2000; Rabbitt, 1966; Surprenant et al., 1999), suggesting that the use of syntactic or semantic context to enhance speech intelligibility also enhances memory (Novick et al., 2005).

We found that task interference effects were strikingly different between clear and intelligibility-matched degraded speech, supporting an essential role for cognitive control at even the mildest levels of perceptual difficulty. These findings echo reports from individuals with hearing impairments, that sustained perception of (amplified) speech is cognitive fatiguing. Nearly one in four people fitted with hearing aids report rarely using them, and one in five are neutral about, or dissatisfied with, their hearing aids (McCormack & Fortnum, 2013). The “listening effort” that is required to understand speech through hearing aids may be an important reason for this lack of enthusiasm. Our results demonstrate that even minor distractions during perception (i.e., tracking a single target) disrupts processing of mildly degraded speech: this illustrates the need to consider cognitive load when assessing and accommodating listeners with hearing impairment.

## Data and Code Availability

All data and code are available upon request.

